# Direct induction of human neurons from fibroblasts carrying the neuropsychiatric 22q11.2 microdeletion reveals transcriptome- and epigenome-wide alterations

**DOI:** 10.1101/2021.10.14.464344

**Authors:** Carolin Purmann, Cheen Euong Ang, Koji Tanabe, Yue Zhang, Soumya Kundu, Tamas Danko, Shining Ma, Alexis Mitelpunkt, Wing Hung Wong, Jonathan Bernstein, Joachim Hallmayer, Bruce Aronow, Thomas C Südhof, Anshul Kundaje, Marius Wernig, Alexander E. Urban

**Author notes:** These authors contributed equally (Carolin Purmann, Cheen Euong Ang, Koji Tanabe).

## Abstract

Standard methods for the creation of neuronal cells via direct induction from primary tissue use perinatal fibroblasts, which hinders the important study of patient specific genetic lesions such as those underlying neuropsychiatric disorders. To address this we developed a novel method for the direct induction of neuronal cells (induced neuronal cells, iN cells) from adult human fibroblast cells. Reprogramming fibroblasts into iN cells via recombinant virus resulted in cells that stain for markers such as MAP2 and PSA-NCAM and exhibit electrophysiological properties such as action potentials and voltage dependent sodium- and potassium currents that reveal a neuronal phenotype. Transcriptome and chromatin analysis using RNA-Seq, microRNA-Seq and ATAC-Seq, respectively, further confirm neuronal character. 22q11.2 Deletion-Syndrome (22q11DS) is caused by a large 3 million base-pair heterozygous deletion on human chromosome 22 and is strongly associated with neurodevelopmental, neuropsychiatric phenotypes such as schizophrenia and autism. We leverage the direct-iN cell model for the study of genetic neurodevelopmental conditions by presenting gene-by-gene as well as network-wide effects of the 22q11DS deletion on gene expression in human neuronal cells, on several levels of functional genomics analysis. Some of the genes within the 22q11DS deletion boundary exhibit unexpected cell-type-specific changes in transcript levels, and genome-wide we can detect dysregulation of calcium channel subunit genes and other genes known to be involved in autism or schizophrenia, such as NRXN1, as well synaptic pathways. This genome-wide effect on gene expression can also be observed at the microRNA and chromatin levels, showing that the iN cells have indeed converted to a neuronal phenotype at several regulatory levels: chromatin, protein-coding RNAs and microRNAs, revealing relevant disease pathways and genes. We present this model of inducing neurons from fibroblasts as a useful general resource to study the genetic and molecular basis of normal and abnormal brain development and brain function.

## Introduction

Many of the major neuropsychiatric disorders such as schizophrenia, bipolar disorder and the autism spectrum disorder (ASD) have a strong genetic predisposition and often a neurodevelopmental basis. Genetic candidate loci for these disorders have recently emerged in considerable numbers^1,2,3,4,5^. Several of these loci consist of complex rearrangements of the genome such as large copy number variants (CNVs) that are affecting multiple genes at once. This poses a challenge for understanding the molecular and cellular mechanisms that are connecting genotype to phenotype, but also offer a promising point of entry into these disease’s etiologies^6^. Many of the genes within these CNVs may have disease-relevant roles that are cell-type specific, dependent on the developmental time point, part of a complex network of interaction with genes outside the CNV, or a combination of these factors. This highlights the importance of studying the effects of neuropsychiatric genetic variants in carefully chosen tissue samples or, absent such samples, tissue culture models of high physiological relevance. There is of course great difficulty acquiring primary brain samples from neuropsychiatric patients. And it is virtually impossible to acquire brain samples that both contain live cells amenable to molecular and neurophysiological analysis while also coming from individuals carrying a relevant genetic variant. In this context, one very promising avenue is the use of induced pluripotent stem cells (iPSCs) to generate neuronal model systems carrying known genetic lesions^7,8^. But the iPSC approach is also not entirely without limitations. For example there is an extended period of time between acquiring the initial tissue sample from a donor and having neuronal cells in culture ready for analysis, which includes upwards of three months for generating the iPSC lines and a further two months or more until functional neurons can be studied. Over this extended period of time cell lines from different patients and controls may begin diverging in their cellular characteristics, making it more challenging to carry out controlled experiments such as taking functional genomics or electrophysiological measurements. In addition to the long development time, iPS cell lines are technically challenging and expensive, and therefore difficult to scale up in large studies.

Here we present a novel cell reprogramming system that allows for the induction of functional human neuronal cells directly from adult fibroblast cells. Importantly these fibroblasts do not have to be from fetal or perinatal samples, as was required for most earlier protocols^9,10,11^, and the resulting induced neuronal cells (iN cells) are of a quantity and quality that allows for multi-level functional genomics and epigenomics analysis of the global molecular consequences of a given aberration in the genome sequence. We demonstrate this and validate the iN cell approach by generating iN cells from donors with the neuropsychiatric microdeletion on chromosome 22q11.2 and then carrying out comprehensive analyses of gene and micro-RNA expression levels as well as of chromatin states, on chromosome 22q and genome wide. We detect cell type specific alterations in the molecular networks controlling gene activity that open new avenues of understanding the molecular etiology of 22q11-Deletion-Syndrome (22q11DS) and its component disorders.

We had previously reported the generation of induced neurons directly from mouse embryonic fibroblasts^11^, mouse postnatal fibroblasts^9^, and human fetal and postnatal fibroblasts^10^. The novel direct induction approach reported here is optimized in several ways and allows, importantly, the use of fibroblast cells from adult donors as starting point. This opens up multiple new avenues of application for fibroblast iN cells, such as the use of donor cells that carry known neuropsychiatric genetic variants and for which the associated clinical phenotypes can be obtained.

At the core of the new direct conversion method is a combination of retroviral induction and a combination of small molecules. The generated neurons are functional and exhibit voltage-gated sodium and potassium currents as well as action potentials. To determine the utility of iNs generated by this novel approach we generated neuronal cells from a cohort of patients with 22q11DS. This developmental syndrome is caused by the most common large-scale chromosomal aberration after Down syndrome, a heterozygous deletion of 3 million basepairs on chromosome 22q in region 11.2. This 22q11.2 deletion is the genetic variant with the strongest known association with schizophrenia and also confers a high risk of ASD^12^.The large deletion CNV encompasses around 50 genes and is strongly suspected of having effects on multiple levels of molecular control and function that will require detailed study in highly physiologically relevant model systems^13^. Currently, despite over twenty years of research into 22q11DS, the disease mechanism is still far from understood. At the same time the 22q11 deletion is an important area of study not only to understand the syndrome in its own right but also as a promising point of entry for the analysis of the molecular basis of major public health concerns such as schizophrenia and ASD in general.

In this study, we have induced neurophysiologically active neurons from adult human dermal fibroblasts using a new strategy based on a combinatorial chemical screen. The neurons fire action potentials and show voltage-dependent sodium and potassium currents. Additionally, transcriptome analysis confirmed conversion into neuronal cell types. When generating neuronal cultures from control probands and patients with a 22q11.2 deletion, we observed downregulation of gene expression in the deletion region both in fibroblasts and neurons, as well as transcriptome-wide, cell-type specific changes in gene expression patterns. The analysis of genome-wide expression patterns in neurons carrying the deletion compared to healthy controls revealed changes in a number of synaptic pathways, especially in postsynaptic components and in voltage-gated calcium channel genes. iN cells generated from 22q11DS fibroblasts also show distinct chromatin and microRNA profiles. MicroRNAs that are dysregulated in 22q11DS target mRNAs that converge on neuron-specific functions. Taken together, we observe local as well as global effects on essential levels of epigenomic regulation of gene activity via chromatin states and miRNA levels. This demonstrates the utility of the directly converted iN cell model for the study of genetic neurodevelopmental conditions while also giving guidance towards the further analysis of 22q11DS.

## Results

### Induction of neurons from patient and control dermal fibroblasts demonstrated by surface markers, electrophysiology and gene expression profiling

As a first step towards improving the reprogramming of human adult fibroblasts to neurons we screened a collection of well-characterized small molecules targeting key developmental pathways individually and in pairs and examined the reprogramming efficiency two and a half weeks after infection in three independent replicates. This effort identified a subset of small molecules that could improve the reprogramming efficiency by up to two-fold (Fig. 1b). The top group of molecules was then tested independently in all possible 2-compound combinations (Fig. 1c). These data showed that molecules modulating the BMP-, TGF-β, Retinoic acid-, and Notch-pathways are represented in the pairs that facilitate neuronal induction. Similar to what has been observed for neural differentiation of stem cells, reprogramming efficiency was greatly improved by the combined blockade of BMP and TGFβ-pathways together with Forskolin, a compound stimulating the enzyme adenylyl cyclase, thus stabilizing and increasing intracellular cAMP levels (Fig. 1e, f). In conclusion, after testing several small molecules and their combinations, we selected the combination of Forskolin, Dorsomorphin, and SB431542 for the induction of neurons from fibroblasts.

**Figure 1:**
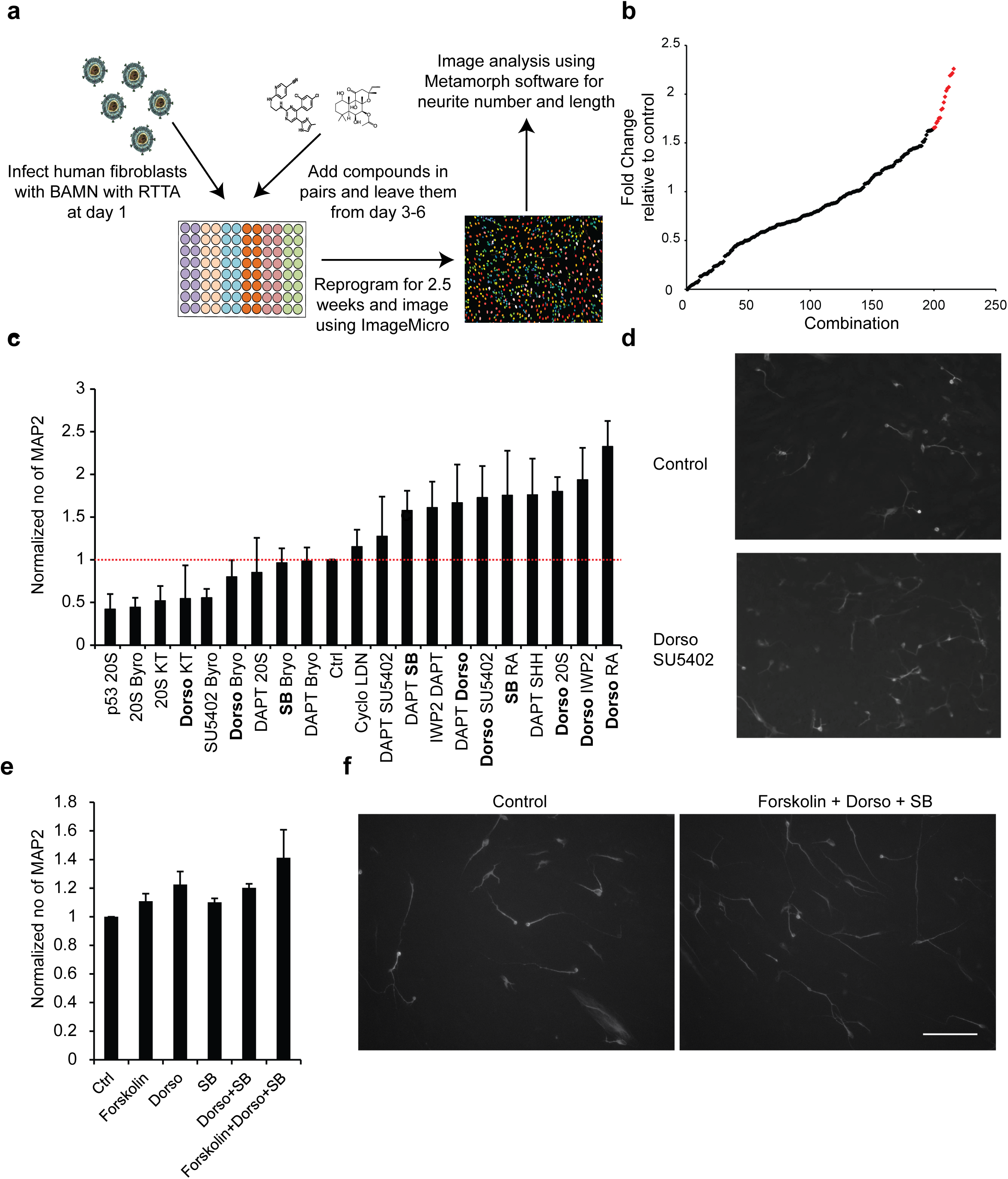
Optimizing the environmental stimuli to convert fibroblasts to iN cells. (**a**) Schematic showing the small molecule screening strategy. **(b**) The foldchange in reprogramming efficiency for various combinations of two drugs. Red: small-molecule combinations that led to >1.5-fold improvement in reprogramming efficiency over control. (**c**) Validations of the drug combinations that gave more than 1.5-fold increase in reprogramming efficiency as measured in MAP2 stained cells. The dual SMAD inhibitors (TGFβ and BMP) are in bold. (**d**) Increased MAP2 staining after treatment with dorsomorphin and SU5402 demonstrates improvement of reprogramming efficiency. (**e, f**) Normalized number of MAP2 positive iN cells with different combinations of Forskolin (PKA activation), Dorsomorphin (BMP inhibition) and SB431542 (TGFβ inhibition), and exemplary micrograph of incrased MAP2 staining after application of small-molecule cocktail.

We next sought to optimize the delivery of the reprogramming factors. Due to efficient gene transfer, lentiviral delivery has been used routinely for reprogramming purposes. Nevertheless, experience in iPS cell reprogramming has shown that absolute expression levels of the reprogramming factors may not be critical and in particular, Moloney Murine Leukemia virus-based vectors showed consistently high reprogramming efficiency^14^. We therefore tested whether oncoretrovirus-based delivery of the four neuronal reprogramming factors Brn2, Ascl1, Myt1l, and Ngn2 (BAMN) would be a more optimal way to reprogram cells (Fig. 2a). Remarkably, by changing the delivery method to oncoretroviral vectors, we were able to observe cells displaying obvious neuronal morphology as early as 7 days after the start of the induction process from adult fibroblasts (Fig. 2b). This observation was striking because using our traditional lentiviral delivery, neuronal cells typically appear only 10-14 days after infection^10^. At 21 days neuronal cells induced by oncoretroviral delivery expressed the panneuronal markers PSA-NCAM and MAP2 (Fig. 2c). In addition to accelerating the reprogramming process, the new approach also improved the reprogramming efficiency based on quantifying the number of MAP2-positive cells on day 21 (Fig. 2 d-f). Importantly, combining the retroviral protocol with the three small molecules (3 sm), i. e. dual SMAD inhibition and Forskolin, improved the reprogramming efficiency even further (Fig. 2g, h). Electrophysiological characterization showed that iN cells generated using this improved protocol fire repetitive action potentials demonstrating neuronal function (Fig. 2 i-k).

**Figure 2:**
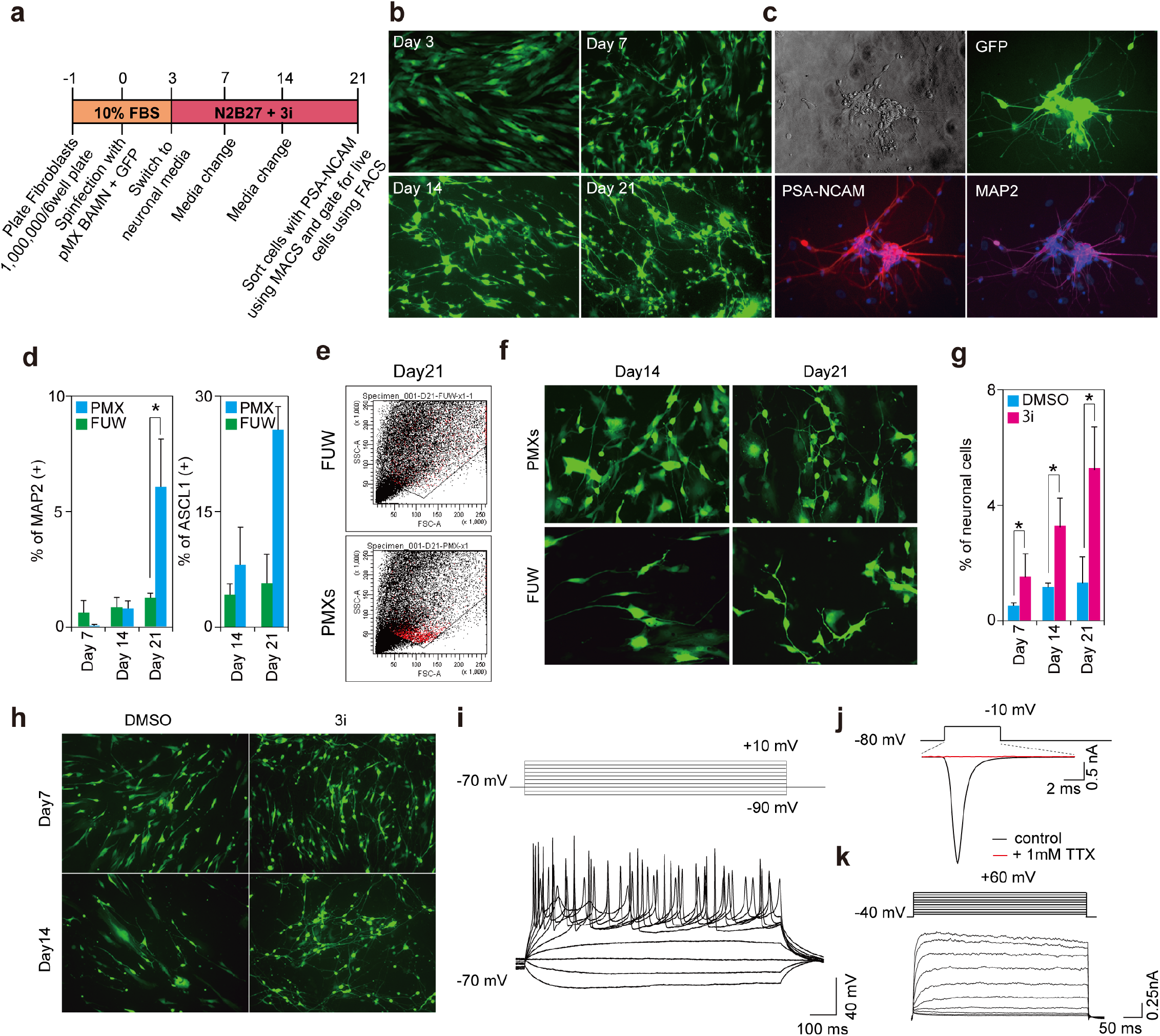
Optimizing retroviral delivery of the reprogramming factors for conversion of fibroblasts to iN cells. (**a**) Schematic of the improved iN reprogramming protocol. (**b**) Progression of iN generation with retroviral delivery of BRN2, NGN2-T2A-ASCL1 and MYT1L at days 3, 7, 14 and 21. Induced neuronal cells appear as early as day 7. (**c**) Bright field, GFP, PSA-NCAM and MAP2 staining of 21 iN cells on day 21. (**d**) Comparison of the percentage of MAP2-positive and ASCL1-positive cells at day 14 and 21 after PMX (retroviral) or FUW (lentiviral) delivery of reprogramming factors. (**e**) FACS plot showing the size of distribution of day 21 iN cells after delivery by PMX or FUW. iN cells generated using the PMX vector are more consistent in size. (**f**) Comparison of iN cells on day 14 and 21 generated by the PMX or FUW delivery systems. (**g, h**) Percentage and visual abundance by GFP of iN cells generated using the PMX system at days 7, 14 and 21, treated with DMSO or with 3i cocktail (Forskolin, SB431542 or Dorsomorphin). (**i**) Sample traces of the multiple action potentials exhibited by iN cells at 4-5 weeks generated by a step-current injection. (**j**) Sample traces of voltage-gated sodium and potassium currents from iN cells generated with the PMX system and addition of 3i.

Next, we applied this optimized neuronal induction protocol to fibroblasts from three 22q11DS cases and three healthy controls from the same collection of biopsy samples with donor age ranging from 1 to 28 years (Fig. S1a). Whole genome sequencing confirmed the 22q11.2 deletion in all three cases (Fig. S1b). Following our optimized protocol we were able to derive iN cell cultures from all six fibroblast lines irrespective of donor age (Fig. S1c, d). There was some morphological variability but no overt and consistent morphological differences between control or deletion iN cell population, as shown by epifluorescence microscopy after ubiquitous labelling of the cells with GFP (Fig. S1d).

We then measured overall gene expression levels by RNA-Seq of both the donor fibroblasts and their iN cell derivatives. Differential expression analysis showed that sets of genes that are statistically significantly enriched in the iN cell transcriptome, as defined by gene ontology (GO) terms, compared to the fibroblast signature are almost all related to neuronal processes (Fig. S2a). In order to assess the iN cell-specific gene expression changes in an unbiased way, we summarized the identified 83 GO terms that are upregulated in iN cells compared to fibroblasts using REVIGO^15^. Three representative cellular components were identified: synapse, synaptic vesicle, and voltage-gated calcium channel complex, which strongly indicates a neural fate of these cells (Fig. S2b). We then searched the RNA-Seq data for expression signatures of neuronal subtypes. As we reported earlier for iN cells generated from perinatal fibroblasts^9,10^, we found robust expression of several markers of excitatory glutamatergic neurons, and low levels of inhibitory GABAergic and dopaminergic markers (Fig. S2c).

Taken together, these gene expression analyses as well as the cell type specific surface markers and the electrophysiological validation show that we have developed and implemented an optimized cell culture protocol to efficiently generate functional neuronal cell types from adult fibroblasts of both control donors and patients with a known, strong genetic risk factor for neurodevelopmental and neuropsychiatric disorder.

### Induced neurons with the 22q11.2 deletion exhibit local and global changes in gene expression

We carried out multilevel functional genomics analyses of the effects of the 22q11.2 deletion. First, we investigated the consequences of the 22q11.2 deletion on the transcriptome. As expected for a genomic deletion, most genes in the deletion region show a drop in RNA levels (Fig. 3a). The 22q11.2 region shows a higher percentage of genes with statistically significant changes in gene expression when compared to the rest of chromosome 22, the whole genome, and a given list of transcription factor genes (Fig. S3a). We also compared expression of individual genes in both cohorts in fibroblast and iN cells. Remarkably, we observed cell-type specific differential gene expression (Fig. S3b). In addition to genes within the deletion boundaries, we also found 38 differentially expressed genes on chromosome 22q but outside of the CNV, 2231 differentially expressed genes genome-wide. While differentially expressed genes within the 22q11.2 deletion boundaries are almost all down-regulated, differentially expressed genes located outside of the CNV are both up- or down-regulated.

**Figure 3:**
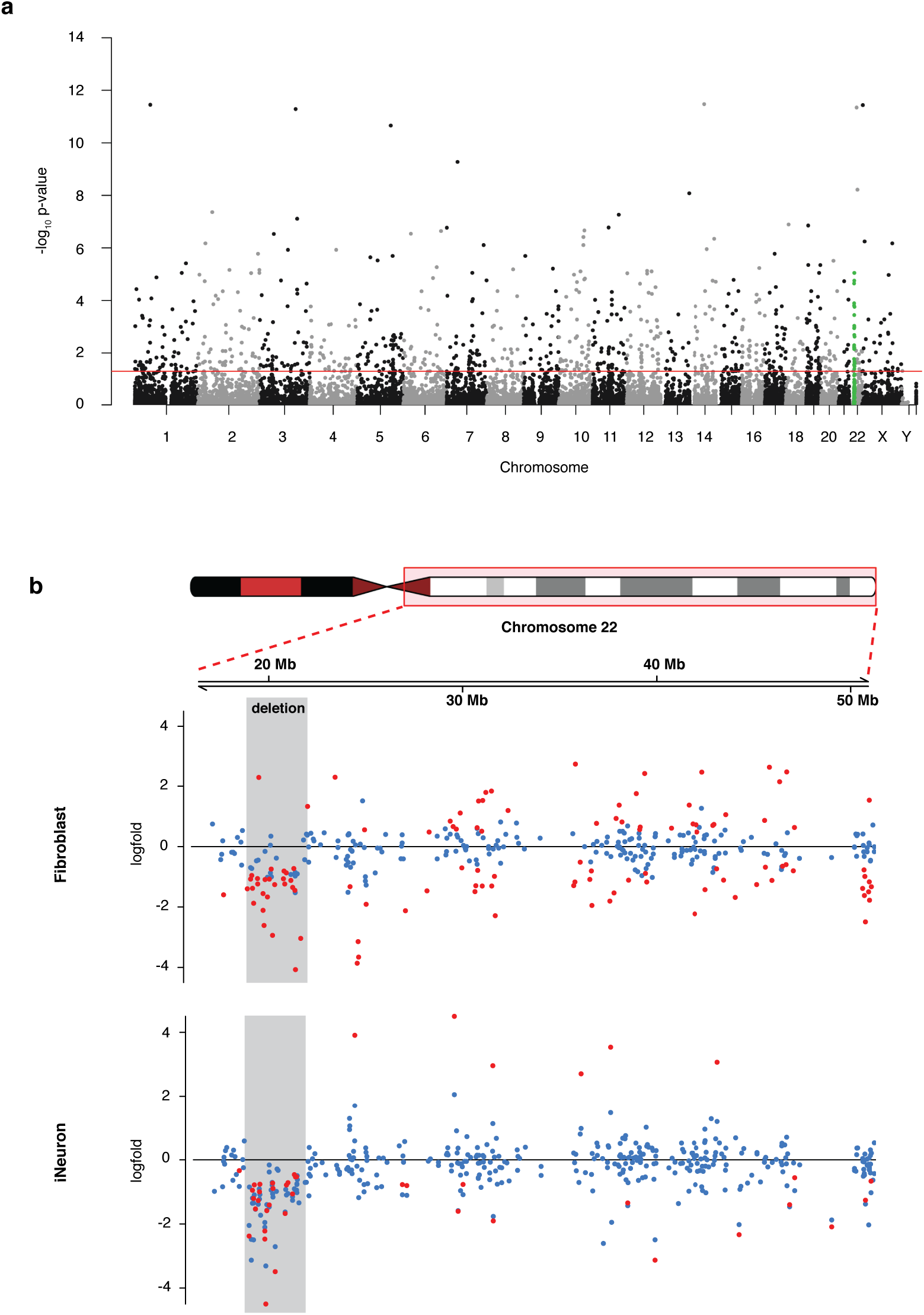
Gene expression in cells with 22q11.2 deletion differs significantly and dependent on differentiation state. (a) Scatter plot showing fold-change of gene expression of the genes on chromosome 22q. Genes are ordered by physical location on the long arm of chromosome 22. The blue box marks the deletion region. Dots in red show a statistical significant fold-change between cases and controls. Blue dots are genes that are not statistically significantly changed. (b) Manhattan plot showing differentially expressed genes in iNs genome-wide. Genes in the 22q11.2 deletion region are plotted in green.

### ATAC-Seq shows globally deregulated chromatin states in neurons with the 22q11.2 deletion

Global changes to gene expression patterns may have an epigenomic underpinning. Earlier we demonstrated that chromatin states are altered globally in lymphoblastoid cell lines (LCLs) with the 22q11.2 deletion^13^, an interesting finding, but in a cell type with limited physiological relevance to the neuropsychiatric phenotypes seen in 22q11DS. We performed ATAC-Seq in the fibroblasts and iN cells, to investigate whether there is a cell-type specific effect of the 22q11.2 deletion in disease-relevant cells, and further to also demonstrate that the novel protocol for iN cell generation allows for such epigenomic analyses.

ATAC-Seq analysis showed that fibroblasts and iN cells have globally different patterns of chromatin openness, and notably control iN cells and 22q11.2 deletion iN cells can be distinguished on the basis of chromatin states alone (Fig. 4a, b). This effect even holds true when carrying out PCA after removing differential chromatin marks located in the 22q11.2 region, and when removing all marks from chromosome 22q (Fig. S4). This underlines the interpretation that there is a genome-wide alteration on the level of chromatin state in neural cells with the 22q11.2 deletion.

**Figure 4:**
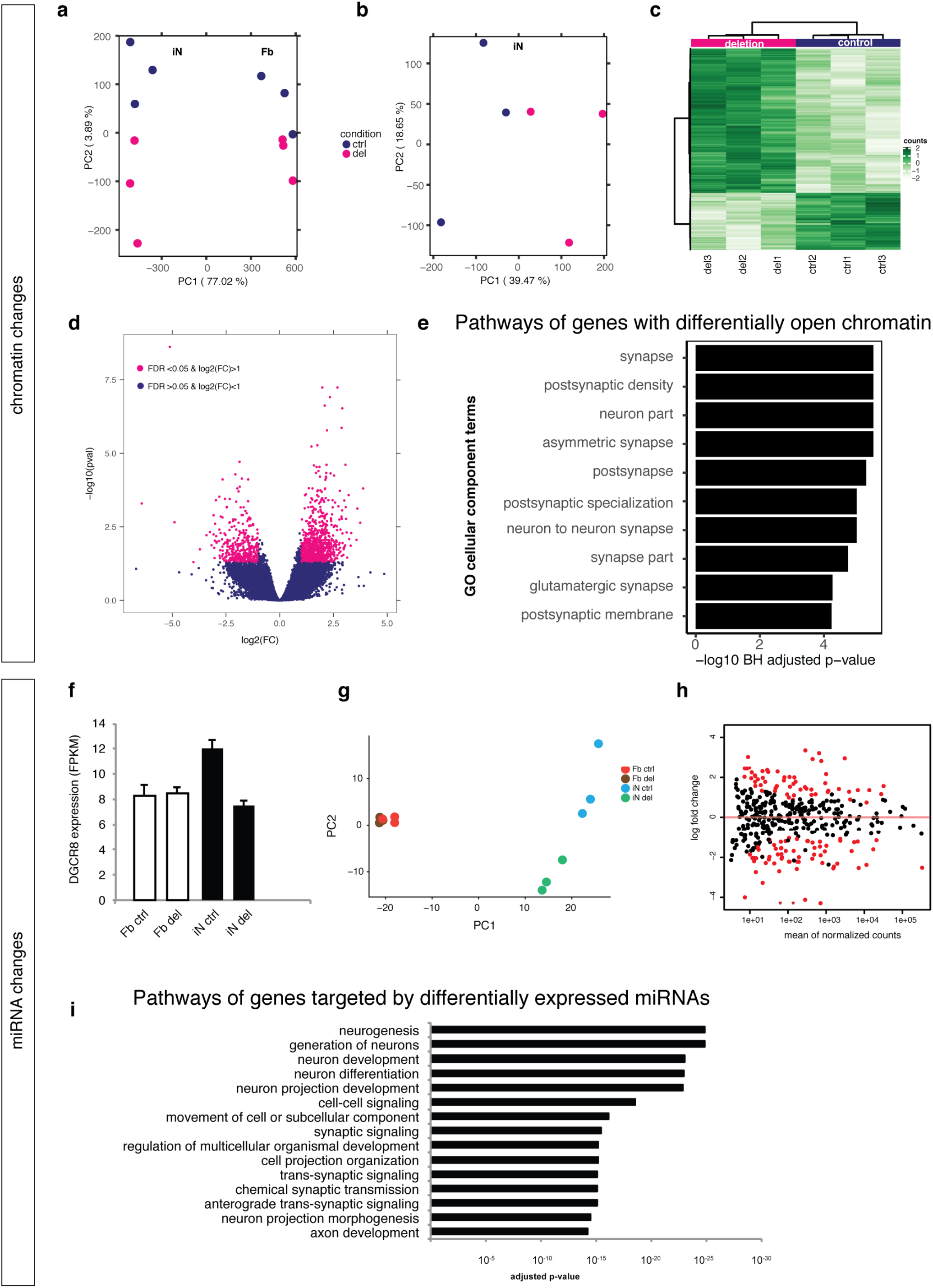
Chromatin and miRNA analysis of reprogrammed iN cells. Principal component analysis of ATAC-Seq signals shows clear difference in chromatin state in (a) fibroblast vs iN, and (b) iN control vs deletion cells. Heatmap (c) and volcano plot (d) of differentially open regions. (e) Pathway analysis of ATAC-Seq signals that differ between deletion and controls iN cells shows enrichment in neural pathways. (f) MicroRNA biogenesis gene DGCR8 is differentially expressed in iNs. (g) Principal component analysis of Fb and iN miRNA expression. There is strong separation between controls and patients in the iN cells. (h) Differential expression of microRNAs in iN cells. Each dot is a micro-RNA (minimum expression level 30 reads). Red dots: microRNAs that are significantly differentially expressed in 22q11.2 deletion iNs relative to control iNs, 102 differentially expressed microRNAs. (i) Pathway analysis of target genes of significantly differentially expressed microRNAs. The 102 differentially expressed microRNAs shown in Panel B have 1132 target genes. Using these target genes as input for Gene Ontology pathway analysis yields multiple pathways associated with development or function of the nervous system.

For the differential ATAC-Seq peaks from the iN cells, a total of 3777 differential peaks (Fig. 4d,e), we assigned the nearest gene and carried out pathway analyses with these genes. This identified 75 pathways with significant differences based on differential chromatin openness patterns between the control and deletion groups. The top 15 most significant pathways are shown in Fig. 4e. The two most significant pathways are nervous system development and regulation of neuron projection development. Most of the other pathways that have significantly different chromatin signatures are also of a neuronal or developmental and cell morphogenesis nature (Supp. Table 1). A summary of the 75 pathways using REVIGO identifies the following umbrella terms: nervous system development, cell projection organization, biological adhesion, cell adhesion, and regulation of Ras protein signal transduction. These results show that on the chromatin level the cells are primed for three-dimensional navigation through the developing cortex even though the cell culture model is twodimensional, showing that the natural cell biology is still reflected in this model.

### iN cells show widespread expression differences of their microRNAs

Next we carried out comprehensive profiling of microRNA expression levels using miRNA-Seq. MicroRNAs are implicated in neuropsychiatric disorders^16,17,1,18,19,20^ and their analysis in the iN cells, in addition to mRNA levels and chromatin state, demonstrates that this cell culture model provides the opportunity to carry out multi-level functional genomics analysis in a highly disease relevant cell type. Furthermore the 22q11.2 deletion region contains the DGCR8 gene, which is an essential component of the microRNA biogenesis pathway^21,22^ and, unlike in fibroblasts, DGCR8 is downregulated in 22q11.2 iN cells (Fig. 4f). We observed distinct expression profiles between the two cell types by PCA. Strikingly, while the principal component analysis of miRNA expression levels does not show separation of control and 22q11.2 deletion fibroblasts, there is clear separation for the iN cells (Fig. 4g). 70% of fibroblast microRNAs have significant differential expression (FDR 0.05, data not shown) relative to the iN cell microRNAs. Neural miRNAs are upregulated, and fibroblast miRNAs are downregulated (Fig. S5). As microRNAs play important roles during development by establishing and maintaining the differentiation of somatic cells, these distinct profiles resulting from before and after direct induction also support the success of reprogramming into iN cells, demonstrating the effects of reprogramming at the level of microRNAs, in addition to the levels of mRNAs and chromatin marks. Between control and 22q11.2 iNs there are 102 microRNAs that are significantly differentially expressed (FDR 0.05, 48 up, 54 down, Fig. 4h). In order to determine the potential functional impacts of these microRNAs we used the Ingenuity Pathway Analysis (IPA) algorithm to predict their target mRNAs. Only targets that are experimentally validated or predicted with high confidence were retained; this yielded 1,132 miRNA target genes that were significantly dysregulated between control and 22q11DS iN cells. Pathway analysis of these target genes yielded 154 GO terms for cellular components, including membrane region, transport vesicle, synapse part, somatodendritic compartment, cell junctions; and 730 GO terms for biological processes including neuron projection development, chemical, synaptic transmission, regulation of cell migration, regulation of neurotransmitter level (the top 15 GO terms are shown in Fig. 4i).

### Network analyses

We carried out two kinds of analyses of the functional networks affected by the changes on the RNA levels. First we used gene set enrichment analysis (GSEA) of the differentially expressed mRNAs, which resulted in 32 significant GO terms (Fig. S6a). Two overlapping GO terms produced the most significant findings: *voltage-gated calcium channel complex* and *voltage-gated calcium channel activity* (FDR q-val of 0.052 and 0.06, respectively). The gene set for *voltagegated calcium channel complex* comprises 15 genes, the gene set for *voltagegated calcium channel activity* comprises the same 15 genes plus three more genes: CACNG4, CACNG5, and CACNA1A (Fig. S6b). None of these genes are within the 22q11.2 deletion region, and they contain single nucleotide polymorphisms that have been associated with risk of schizophrenia, bipolar disorder and of mood disorders^1,23,24^. Most of these 18 genes are down-regulated in iN cells carrying the 22q11.2 deletion (Fig. S6b, c). The deregulation of this entire class of Ca^2+^ channel genes would not have been observed when analyzing fibroblasts from 22q11DS patients, many of these genes are not expressed or are not differentially expressed in fibroblasts (Fig. S6c). As before, we summarized this list of GO terms using REVIGO resulting in four categories: *synapse, synaptic vesicle, voltage-gated calcium channel complex*, and *chromatin remodelling complex*.

To further dissect the changes in RNA expression, we filtered for the significantly changed genes with an at least two-fold expression change. These genes were further sorted into different modules depending on their expression pattern in the two cell types (fibroblast or iN cells) and the two genotypes (control or 22q11 deletion) (Fig. 5). Genes fell into one of eight modules: *Module 1* (188 genes) higher expression in controls in both fibroblasts and iN cells, *Module 2* (342 genes) higher expression in deletion cells in both fibroblasts and iN cells, *Module 3* (473 genes) higher expression in fibroblasts than iN cells in both controls and 22q11del, *Module 4* (218 genes) high expression in only control fibroblasts, *Module 5* (408 genes) higher expression in only 22q11del fibroblasts, *Module 6* (318 genes) higher expression in iN cells than fibroblasts for both controls and 22q11del cells, *Module 7* (459 genes) higher expression in iN cells than fibroblasts but with 22q11del cells showing lower expression than iN control cells, and *Module 8* (200 genes) genes with higher expression in iN cells than fibroblasts but with 22q11del iN cells showing even higher expression than control iN cells.

**Figure 5:**
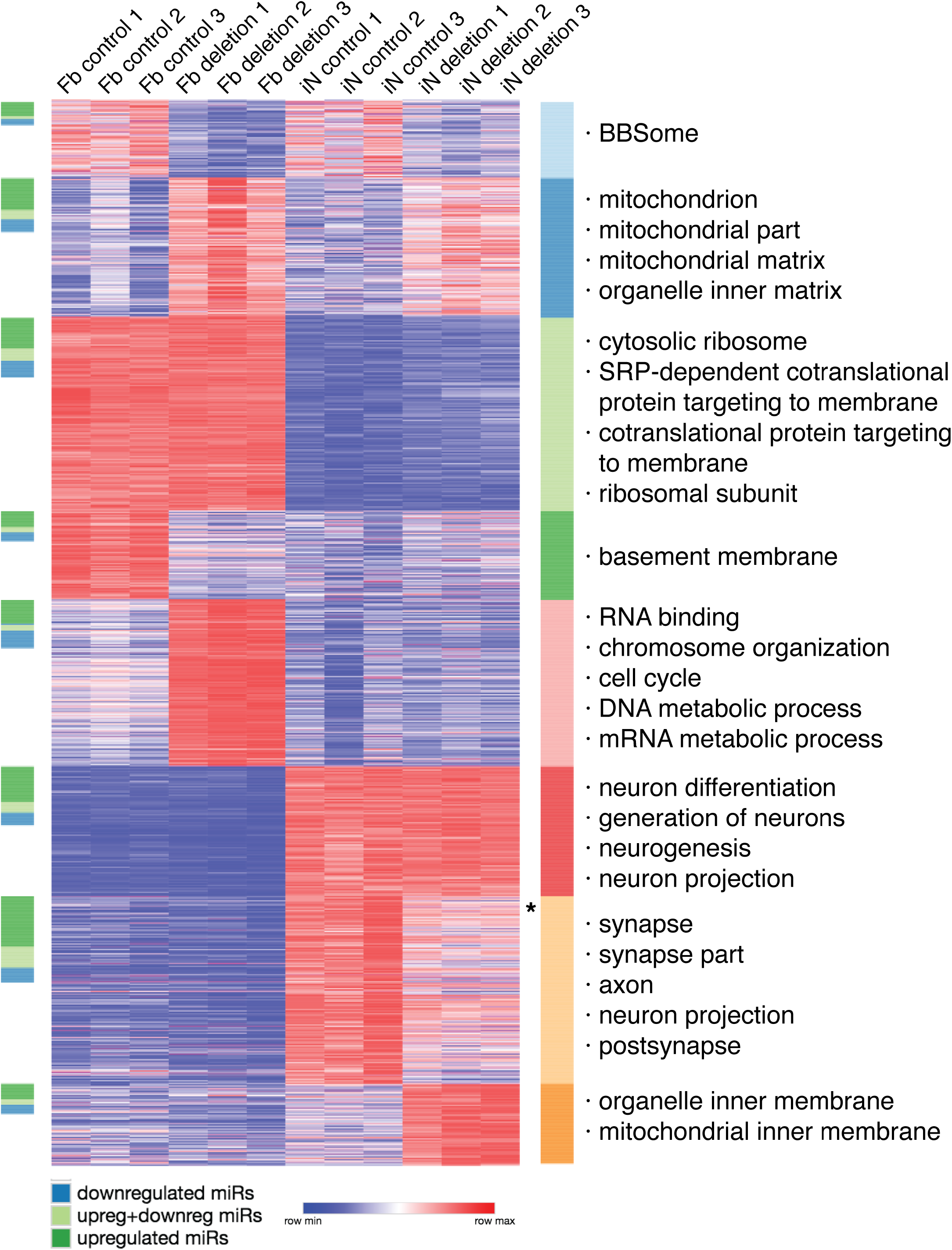
Clustering of differentially expressed 22q11.2 genes and analysis of functional categories of differentially expressed genes. Genes with significantly changed gene expression (>2-fold change) are shown with their relative expression and sorted in rows. Rows are clustered into eight modules depending on their expression pattern in fibroblast and neurons and individual genotype (control or deletion). On the left next to the heatmap a color code (dark green, light green, blue) shows if the corresponding gene in that row is a target of an up- or downregulated miRNA. On the right next to the heatmap, the pathways that are enriched for the genes in each module are listed. Asterisk indicates position of DGCR8 gene in heatmap.

*Module 6*, which comprises genes that are upregulated in iN cells compared to fibroblasts, is enriched for neurogenesis and neuron differentiation. This serves as an additional positive control that the neural induction was successful and that the fibroblasts were converted into neurons.

Genes that were upregulated in 22q11del cells, both in fibroblasts and iN cells, were enriched for mitochondrial pathways. The 22q11 deletion contains a number of mitochondrial genes and a potential involvement of the mitochondria in 22q11DS etiology has been reported^25,26,27^. Particularly interesting is *Module 7*, which contains genes that are upregulated after reprogramming into neurons but where 22q11del cells do not upregulate as much as control cells. The pathway analysis for this module yielded pathways potentially relevant to the neuropsychiatric phenotype of 22q11.2 patients. And also, in this module the genes that are miRNA targets are not evenly distributed between miRNAs that are up- or downregulated in 22q11del iN cells. The pathways that are enriched for genes that are iN-specific but which show lower expression in 22q11del iN cells are synapse, postsynapse, axon, and neuron projection. The genes in this module are more highly enriched for targets of miRNAs that were upregulated in the expression analysis of miRNA-seq of control versus patient iN cells. This is consistent with the expectation that mRNAs become downregulated if they are targets of upregulated miRNAs.

To further compare all pathways enriched in each module, we mapped all pathways enriched for each module (Fig. 6). Visualizing the large number of significant pathways in this unbiased fashion revealed an interesting pattern. Generally, there is very little overlap between pathways that are enriched in each module. The only two exceptions are a small overlap of mitochondrial related pathways in both del-up and iN del-up modules, and a significant larger overlap between iN and iN_del-down modules. Both of the latter two modules contain neuron-related pathways. To tease out the misfunction specifically in 22q11del neurons as opposed to functional differences between fibroblasts and neurons in general, we compared the pathways and noticed that in particular the synaptic and especially the postsynaptic pathways are enriched and also with a more significant p-value (Fig. S7). We plotted the gene expression of well established components of the synapse^28^ (Fig. 7). This highlighted a general trend of downregulation, including of neurexins and their neuroligin binding partners (Fig. S8). The genes that are part of the synapse are enriched for differentially expressed genes (p-value p<0.0001, two-tailed Chi-square test).

**Figure 6:**
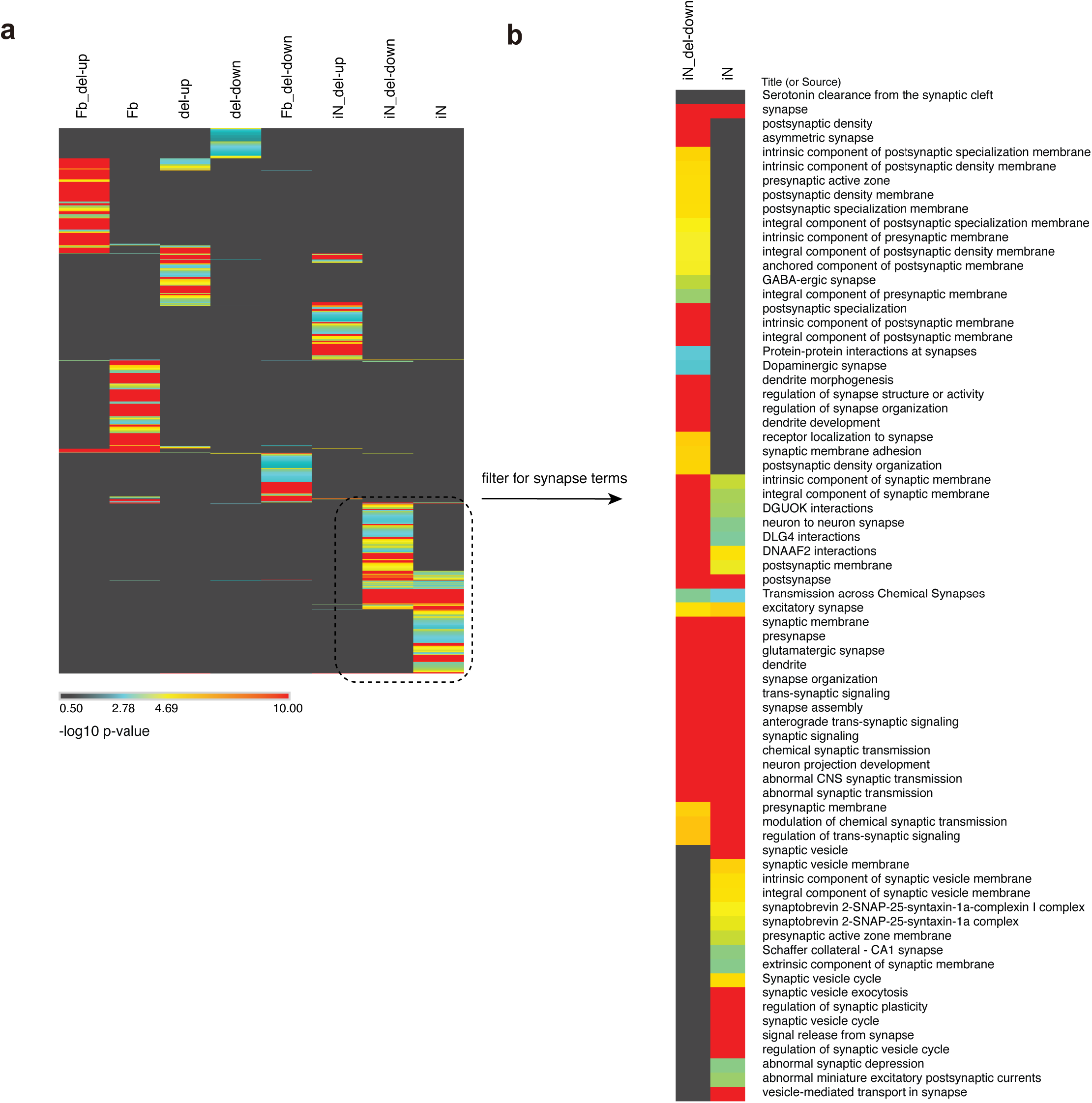
Comparative pathway analysis. (a) Pathway analysis of the eight modules reveals overlap of iN and iN_del-down pathways. (b) By filtering for synaptic pathways it is revealed that synaptic pathways and particularly postsynaptic pathways are more strongly enriched in iN del-down genes (by p-value).

**Figure 7:**
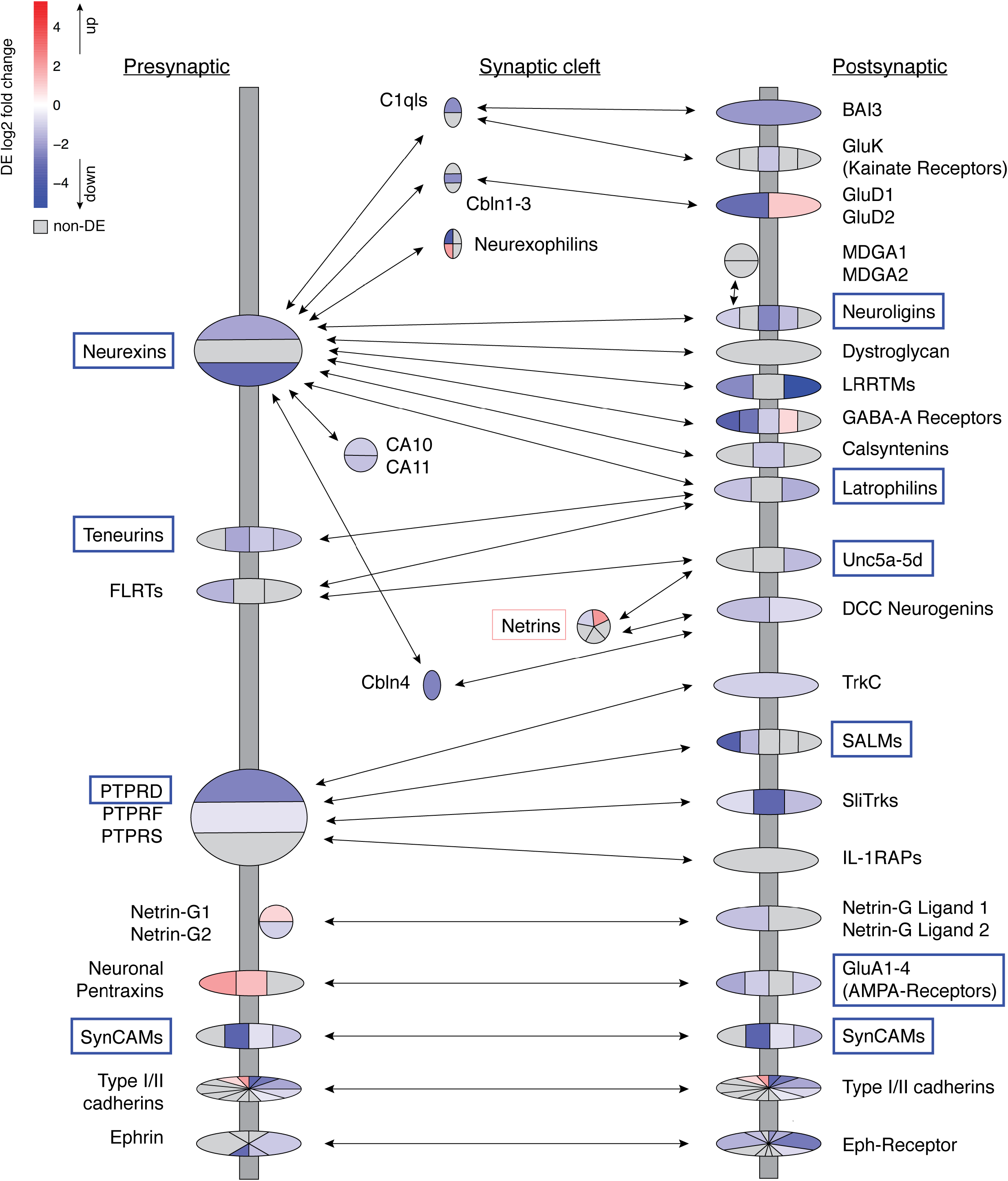
Differential expression in 22q11.2 deletion patient neuronal cells of voltage-regulated calcium channel genes and of genes that form the synapse. Differential levels of gene expression are shown, red is up in patient iNs, blue is down in patient iNs, gray genes are not differentially expressed in iNs. Genes that form the synapse are enriched for downregulated genes. Genes in boxes are targets of differentially expressed miRNAs.

## Discussion

To be able to go from patient to cellular phenotype in the study of psychiatric diseases has been a central aim for neurogenetics and genomics. Our study provides a new model that allows the study of live human neurons from patients with a known genetic lesion and neuropsychiatric phenotype. As a proof-of-principle study we employed our novel and optimized method to generate and study neurons from patient fibroblasts with 22q11.2DS and healthy controls.

The iN cells generated express neuronal markers on RNA and protein levels and possess electrophysiological properties of neurons. The main increase in efficiency was achieved by changing the gene delivery from lentiviral to onco-retroviral vectors. This was a surprising result, since lentiviral infection yields higher expression levels than onco-retroviruses. The infection rate is predicted to be higher because onco-retroviruses can only infect dividing cells, whereas lentiviruses are independent of the cell cycle. We speculate that potentially the expression dynamics may be different between the two viral systems. We have observed in the past that lentiviral expression of one of the key transcription factors, Ascl1, is silenced within 5 days after infection. Ascl1 is oscillating in infected fibroblasts in non-uniform patterns with an average higher expression in cells that move on to become neurons than in cells that fail to reprogram^29^. Onco-retroviral delivery could lead to lower initial but more consistent long term expression, which could lead to an overall increase of reprogramming factor abundance in the time course of reprogramming, and thus improve conversion efficiencies.

Transcriptomic analysis showed a reduction in expression for genes in the deletion region as expected for the 22q11.2DS patients. An analysis of genomewide expression patterns showed downregulation of genes in synaptic pathways, especially post-synaptic genes, chromatin remodelling, and voltage-gated calcium channel complex. Pathway analysis revealed changes in synaptic signalling pathways and axon guidance. MicroRNAs in induced neurons are differentially regulated between deletion and control cells target genes that are disease relevant.

22q11.2DS is a prominent model for neuropsychiatric diseases. Individuals with deletions in 22q11.2 (seen in about 1 in 4000 live births) have a high risk of schizophrenia: around 30%, i.e. an around 30-fold increase in risk, making this the largest known effect of a single mutation on risk of schizophrenia. Additionally, a high prevalence of attention deficit hyperactivity disorder (ADHD), autism spectrum disorder (ASD), and anxiety disorders has been reported^30^. Most of the patients, such as the three patients included in our study here, carry the same 3 Mb deletion with well-described clinical effects. However, the molecular mechanisms by which the 3 Mb deletion increases risk of schizophrenia or ASD is not known, i.e. which deleted gene or genes are primarily responsible, and whether there are important effects outside of the deletion regions, e.g., due to deleted genes that regulate genes in other regions, in spite of over two decades of intense analysis of the 22q11DS locus.

Having a functional model available that allows the study of a relevant cell type facilitates research into the consequences and potential causal mechanisms of disorders such as schizophrenia and ASD by generating testable hypotheses.

By comparing fibroblast and neural expression patterns, we have shown the importance of establishing a neuronal cell type when studying neuropsychiatric disorders with genomic techniques such as RNA-Seq. We have observed celltype specific gene expression. This underscores the importance of studying the impact of a genetic risk factor in the relevant cell type. We found that genes of voltage-gated calcium channel family, including CACNA1C, are differentially expressed. CACNA1C has been identified as risk gene genome-wide association studies for schizophrenia and bipolar disorder. Our more direct approach studying expression levels in neurons compared to SNPs in DNA validates our model for generating testable hypotheses in relevant cell types.

Another class of genes that has been implicated in schizophrenia and ASD are genes that are associated with epigenetic and chromatin regulation^31,32,33,34^.

We also observe changes in chromatin remodelling pathway in the gene expression of 22q11DS iN cells as well as when analyzing the actual chromatin itself. Remarkably, we have also found that patients and controls can be clustered based solely on genome-wide chromatin conformation analysis using ATAC-Seq methods, which means that the chromatin changes precipitated by the presence of the 22q11 deletion are not limited to chromosome 22. Using pathway analysis on ATAC-Seq results, we confirmed changes in nervous system development in 22q11DS that we found with our RNA-Seq analysis, thus cross-validating each approach.

We observed changes in gene expression related to the function of the synapse, in particular with the postsynaptic side. A role for synaptic and postsynaptic changes in schizophrenia have been implicated previously by studying genes affected by small de novo mutations^35^. We observed downregulation of multiple synaptic proteins, including NRXN1. NRXN1 is a transmembrane receptor that is involved in the formation of synaptic contacts. Deletions of NRXN1 cause ASD and schizophrenia. A whole exome sequencing study of autism spectrum disorder patients identified NRXN1 as the one gene being involved in schizophrenia, intellectual disability, congenital heart disease, and epilepsy^34^. Interestingly, 22q11 deletion syndrome includes all of these symptoms, even though the NRXN1 gene does not lie in the deletion region. This could point to a convergence of pathways for neuropsychiatric disorders as well as genetic pleiotropy and phenotypic heterogeneity.

Unlike our RNA-Seq analysis, we also found distinct pathways of neuron projection regulation that have profound differences between patient and controls determined by ATAC-seq. This could indicate that chromatin patterns are established that are not yet reflected in transcript levels. Neuron projection dysregulation is particularly interesting as previous studies have implicated altered neuronal connectivity in 22q11DS^36^. Moreover, MRI studies have found both increased and decreased within-network connectivity in 22q11DS patients^37^. Future studies will determine the molecular impact of our findings. Our results suggest that the chromatin signature of patients already contains the imprint of expression disturbances that only come into effect in the context of more mature neurons or in neurons allowed to navigate through the native developing brain. This again emphasizes the importance of studying cells in states as close to their functional equivalent in the human brain as possible and we do not rule out further changes in expression profiles that could be revealed by studying affected brain regions within patients exhibiting schizophrenia, ASD, or other psychiatric disorders with a neurodevelopmental component. However, since this material is far from being accessible, we believe our model represents an important step forward in bridging that gap. By including 22q11DS as a genetic model for this iN cell proof-of-principle study, we were able to validate the results on both directions: using 22q11 validates the iN cell approach and using neurons validates that the results are relevant to the understanding of 22q11.

In summary, our results clearly demonstrate that disease-associated cellular processes can be observed in induced neuronal cells derived directly from patient skin cells. Thus, patient-derived iN cells represent a valuable new model of human neurodevelopmental and neuropsychiatric disease. This is an important proof-of-concept and suggests the promise that new disease mechanisms can be discovered using this approach, some of which may become the basis for potential therapeutic intervention or personalized medicine. The described protocol allows fast and scalable generation of neurons within three weeks without the need for induced pluripotent stem cells delivering cells with a patient-specific mutation and genetic background that can be used for relevant tests.

## Materials and Methods

### Cell line maintenance and characterization

Subjects with 22q11.2 Deletion Syndrome and healthy controls were recruited from an on-going study at Stanford. The study was approved by the Stanford University Institutional Review Board (IRB). Skin punch biopsies were taken from consenting subjects and primary human dermal fibroblast cell lines were established. Fibroblast cells were used at an early passage number and maintained in FibroLife S2 Fibroblast Medium (Lifeline Cell Technology).

Copy number status of the 22q11.2 region was verified using droplet digital PCR. Briefly, genomic DNA was extracted from fibroblasts and a quantitative PCR was done using digital droplet PCR technology (Bio-Rad) comparing dosage of a probe in the 22q11.2 locus and a control locus (TERT).

### *iN Protocol – screen and* generation and retroviral production

Retroviruses were produced using the protocol described. Briefly, 6μg of pMX and 3μg of VSVg were transfected into plat-gp cells using FuGENE transfection reagents (Promega). Viral supernatant was harvested after 48 hours.

### iN cells generation and purification

Dermal fibroblasts obtained three healthy and 22q11.2 patients were cultured in DMEM (Gibco, 12430) with 10% FBS (Invitrogen). To reprogram, 1×10^6^ fibroblasts were seeded in a 6 well plate. The next day, 2ml of retroviral supernatant with all factors in equal proportion (pMX-Ascl1-T2A-Ngn2, pMX-Brn2, pMX-Myt1l and pMX-GFP) was added per well and the plates were spin-infected at 700g for one hour. Viral supernatant was replaced with 10% FBS media after spinfection. At day 3, media was changed to N2B27 media [DMEM/F12, N2, B27 (Gibco) and 1.6ml of Insulin (Sigma)] with 3i [Dorsomorphin (2μM), Forskolin (3μM) and SB431542 (10μM)]. Media change was performed once every 3 days. At day 21, magnetic activated sorting for PSA-NCAM positive cells were performed according to the manufacturer’s protocols (APC conjugated PSA-NCAM and Anti-APC magnetic beads, Miltenyi Biotech) and live cells were isolated from the MAC sorted PSA-NCAM positive population using FACS.

### Electrophysiology

Whole-cell patch-clamp recordings were performed as described previously.^38^

### RNA-Sequencing

Cell pellets were resuspended in Trizol reagent and stored at −80°C until RNA extraction. RNA extraction was performed using a combination of PhenolChloroform phase separation and silica-based spin column purification, including a DNase I treatment, to ensure high-quality DNAse-free RNA. RNA integrity was checked using Agilent Bioanalyzer 2100 instrument and RNA 6000 Pico reagents. Illumina sequencing libraries were prepared using NEBNext rRNA Depletion and Ultra Directional RNA library prep kits (New England Biolabs) following manufacturer’s instructions. Libraries were sequenced on an Illumina NextSeq500 machine at 2×75bp. RNA-seq data was aligned using STAR and differentially expressed transcripts were identified using cufflinks. Pathway analysis was performed using GSEA and ToppGene suite^39, 40^. GO summation was done using REVIGO^15^.

### MicroRNA-seq

MicroRNA-seq libraries were prepared using NEBNext Multiplex Small RNA Library Prep Set for Illumina (New England Biolabs, cat. no. E7300) following manufacturer’s instructions. The miRNA-library peak was size-selected using the Blue Pippin instrument and 3% Agarose Gel Cassettes (Sage Science). Successful miRNA sequencing library preparation was confirmed using Bioanalyzer High Sensitivity chip reagents (Agilent). Libraries were sequenced on an Illumina NextSeq instrument. Reads were adapter-trimmed and quality-filtered using cutadapt, and then sequentially aligned against rRNA, tRNA, and snoRNA reference sets using bowtie. Remaining unmapped reads were aligned against a miRNA database. Multiple mapping reads were equally distributed among all mappable locations. Differential expression analysis was performed using DESeq2.

### ATAC-seq

ATAC-seq was performed on 40,000-50,000 cells following a published protocol (Buenrostro et al., 2013). qPCR was performed to avoid overamplification. Before sequencing, successful tagmentation and ATAC-seq library construction was confirmed by looking for presence of typical ATAC-seq fragment sizes using Agilent Bioanalyzer 2100 instrument and high sensitivity reagents. Libraries were sequenced on Illumina NextSeq500 machine at 2×75bp. ATAC-seq data was aligned using Bowtie2. Duplicate reads and reads mapping to mitochondrial DNA were removed before peak calling with MACS2. Downstream analysis of peaks was done using R packages Diffbind and ChIPpeakAnno.

---

**Figure S1:**
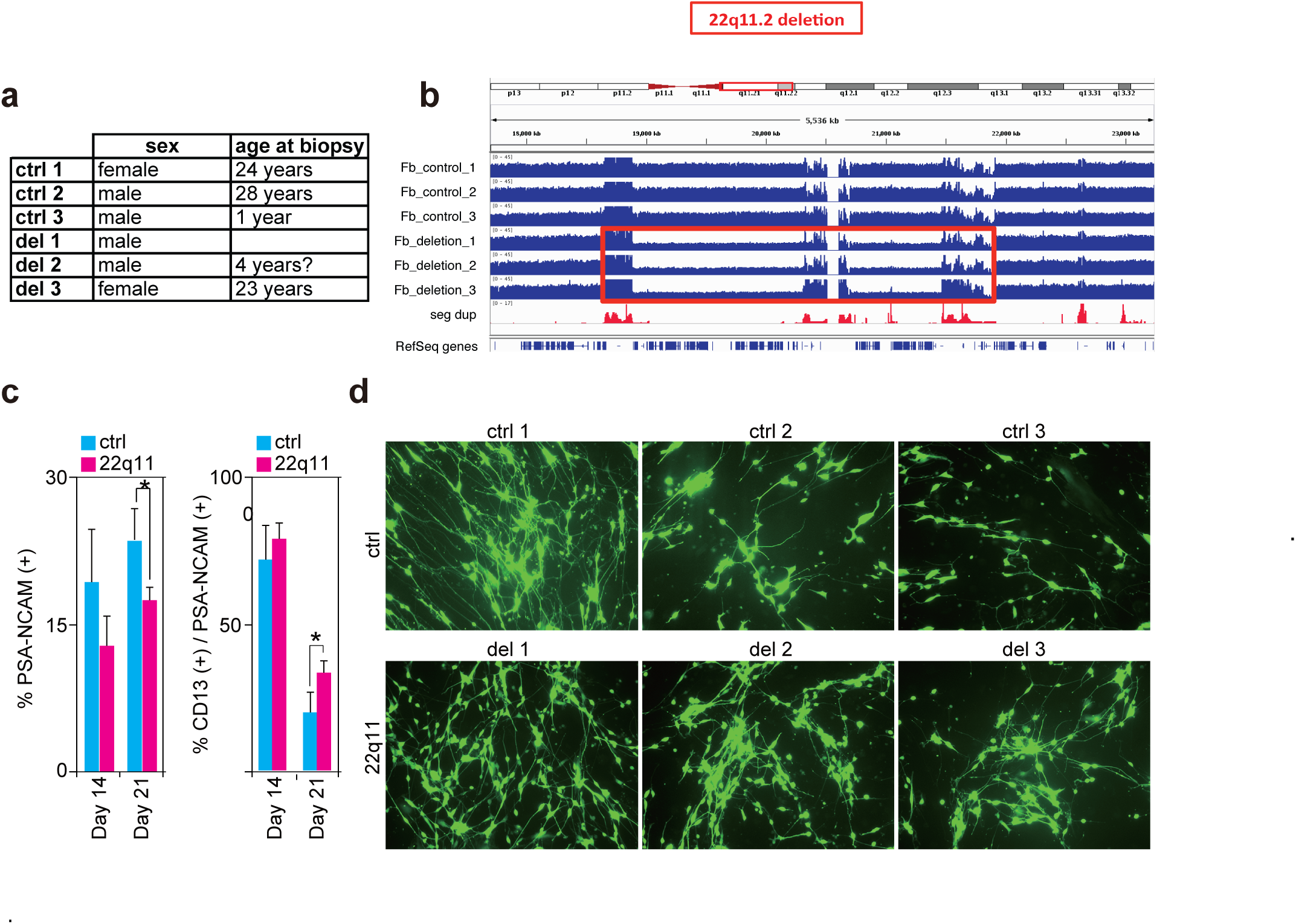
Conversion of control and 22q11.2 deletion syndrome patient fibroblasts to iN cells. (a) Fibroblast cell lines used in this study. (b) Whole genome sequencing of fibroblast DNA. Coverage map shows drop in 22q11.2 region in deletion samples. Noisy regions represent segmental duplication regions. (c) Percentage of PSA-NCAM positive cells in 22q11.2 patients versus controls at day 14 and 21. **(d)** Percentage of CD13 positive / PSA-NCAM positive cells in 22q11.2 patients versus control at day 14 and 21. * represents p<0.05.

**Figure S2:**
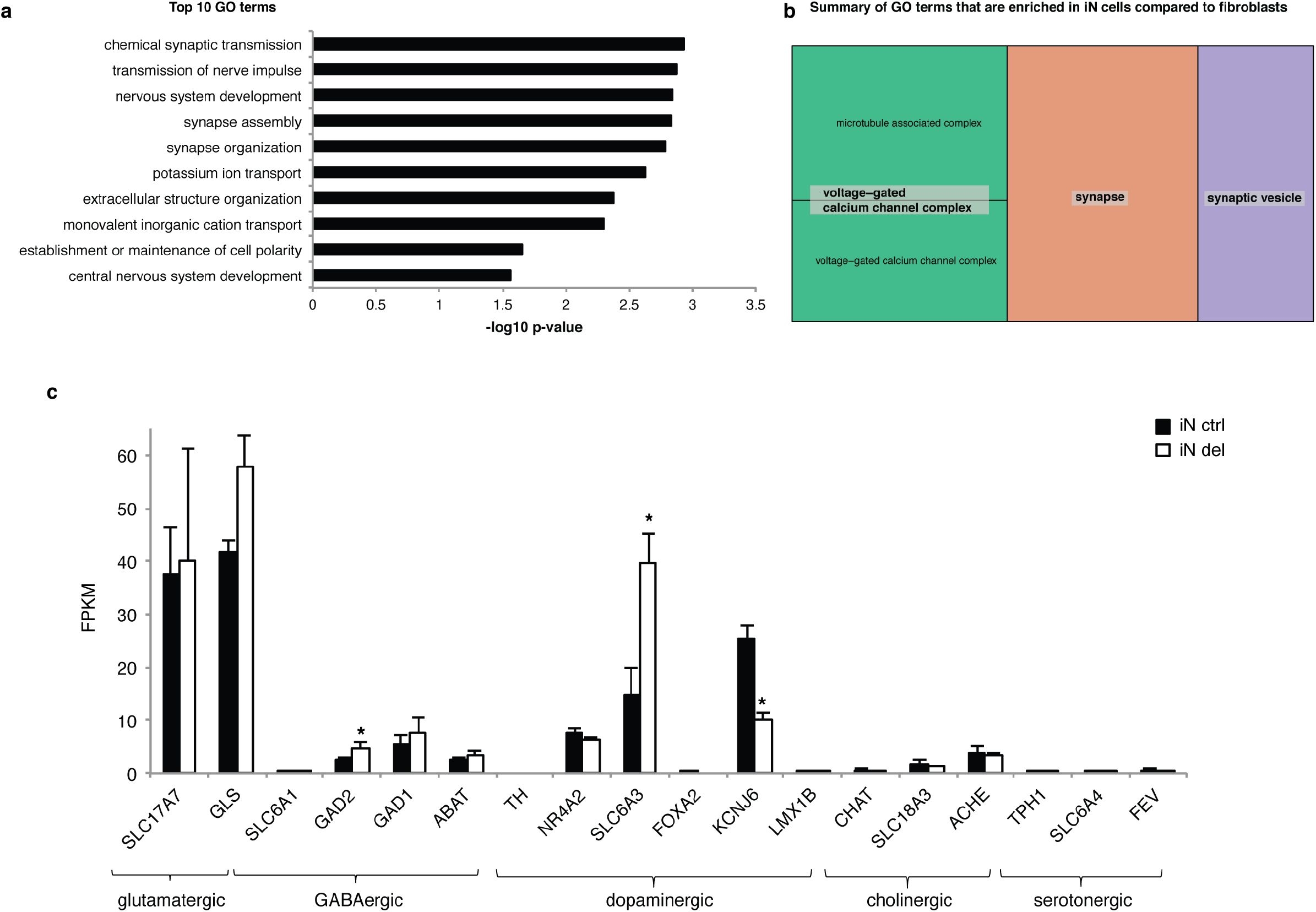
Neurons induced from human dermal fibroblasts show neural expression profile. Bar graphs showing strength of expression of neural marker genes measured by RNA-seq. (a) Bar graph showing p-values for summarized top 10 GO terms that are significantly upregulated in induced neurons versus dermal fibroblasts. (b) REVIGO treemap of GO terms that capture the most difference of Fb ctrl cells vs iN ctrl cells. (c) Bar graph of expression FPKM values (y-axis) of marker genes for different neuronal subtypes (x-axis)

**Figure S3:**
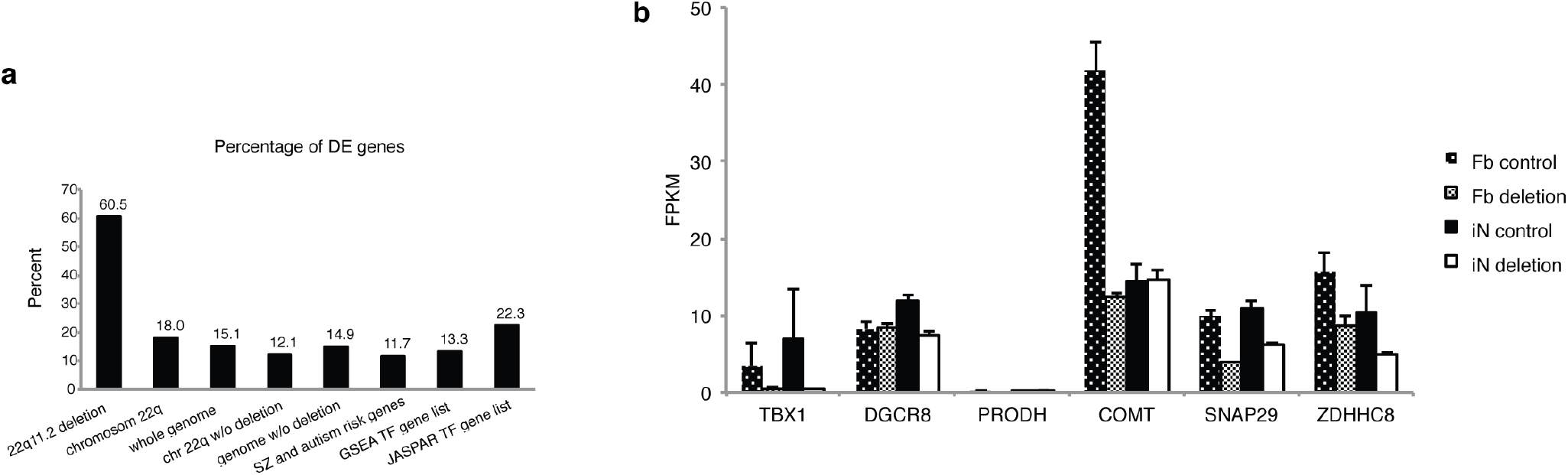
Cell type and genotype specific gene expression changes in reprogrammed iN cells. Percentage of expressed genes that are statistically significantly changed between control and deletion iN cells across different categories of genes. (C) Bar graph showing the expression of genes of interest in the 22q11.2 deletion region. Gene expression in 22q11.2 cells differs significantly dependent on differentiation state.

**Figure S4:**
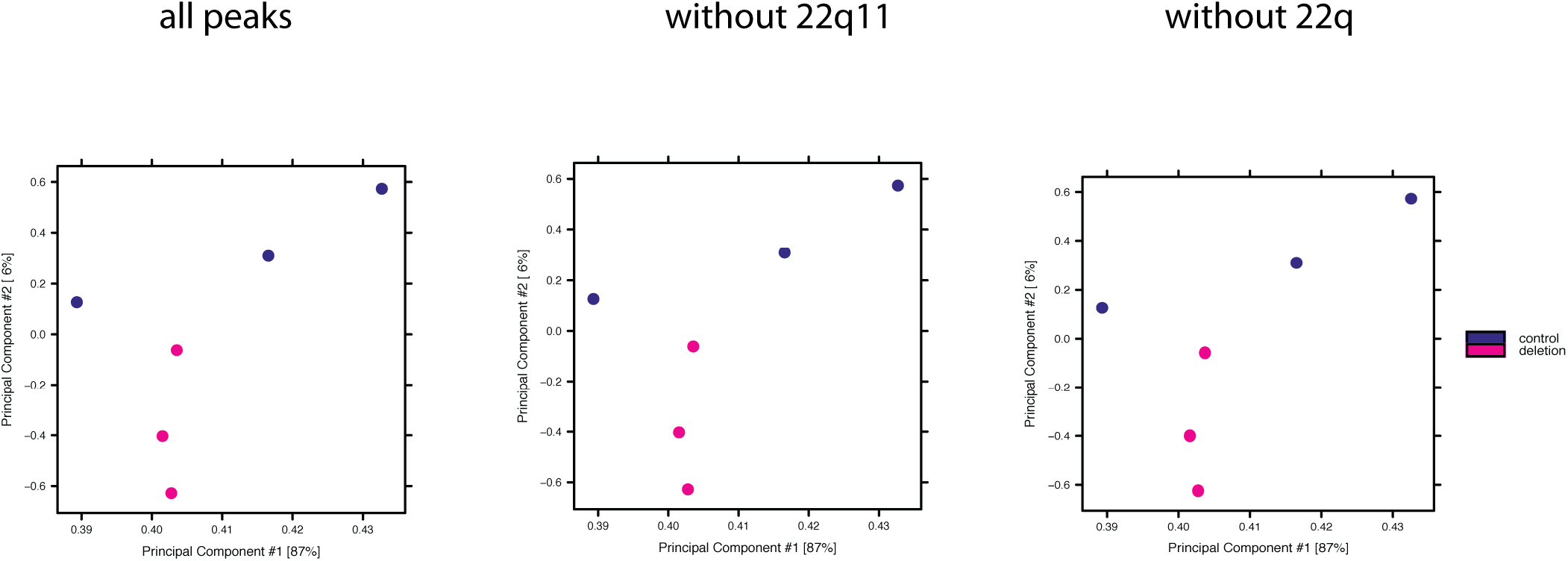
Principal component analysis of open chromatin regions in del and ctrl iN cells. From left to right: PCA plot of regions genome-wide (all peaks), PCA plot of regions excluding the 22q11.2 deletion region (without 22q11), PCA plot of regions excluding all peaks on chromosome 22q (without 22q). Plots are nearly identical showing that separation by genotype is not driven by merely losing one copy of 22q11 genomic material.

**Figure S5:**
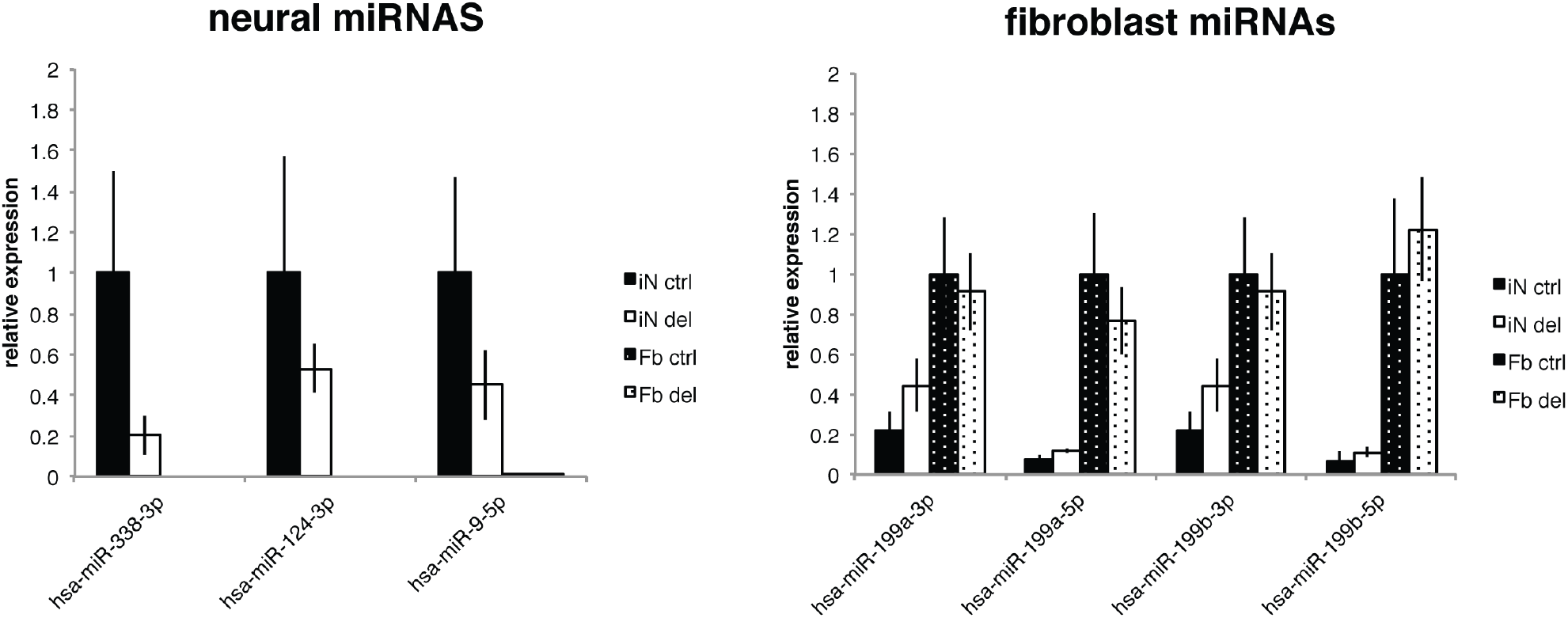
Expression of marker miRNAs in fibroblast and iN cells. Left: Relative expression of neural marker miRNAs^41^, normalized to expression level in iN control cells. Right: Relative expression of fibroblast marker RNAs, normalized to expression level in fibroblast control cells. Fibroblast lines do not express neural miRNAs, but iN cells do. iN cells show significant downregulation of fibroblast miRNAs, indicating a cell type conversion at the miRNA expression level.

**Figure S6:**
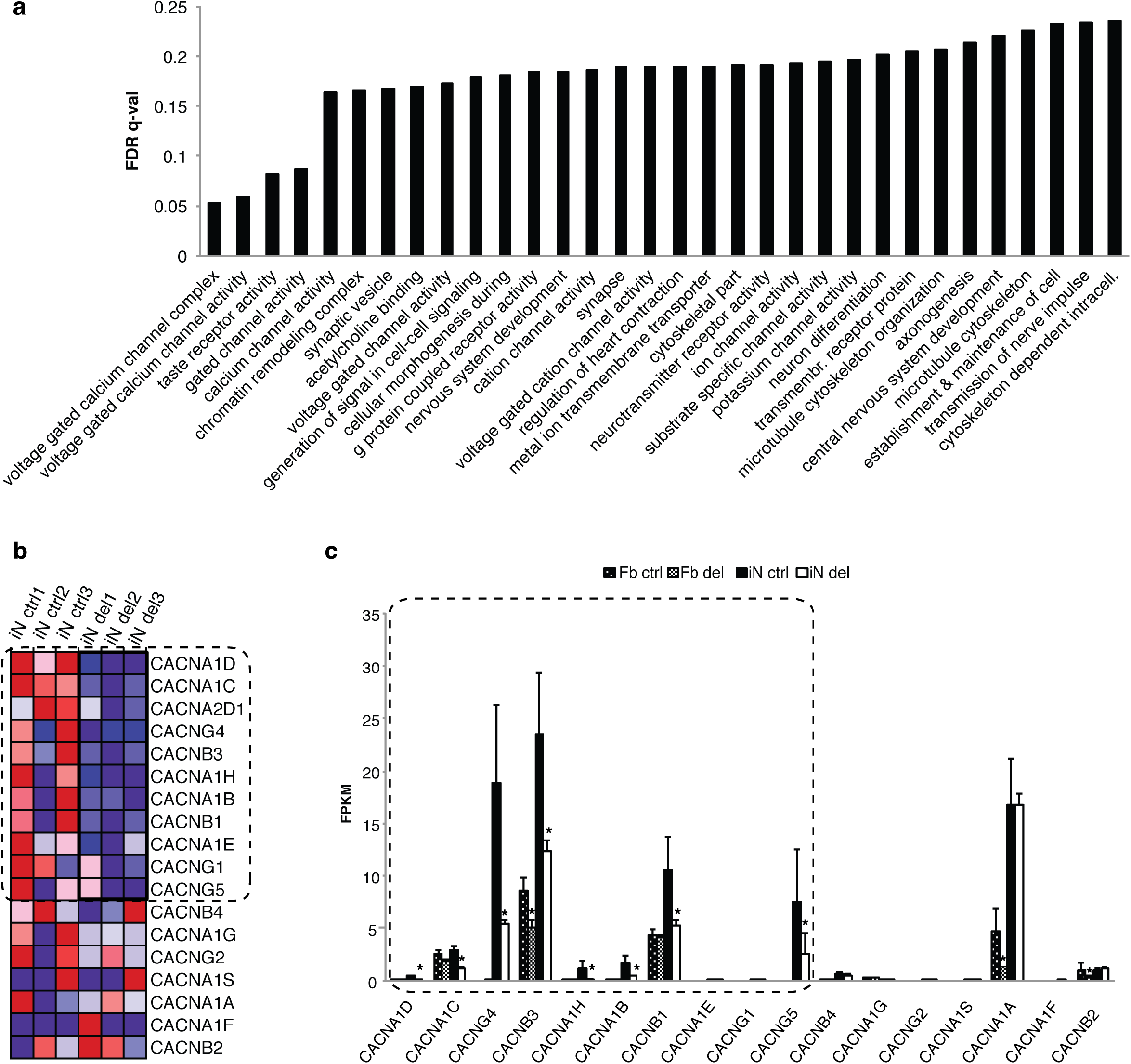
GSEA pathway analysis of iN deletion cells reveals changes in voltage-gated calcium channels. (a) Bar plot showing FDR q-avlues of GSEA pathway analysis. The most significant pathways are *voltage-gated calcium channel complex* and *voltage gated channel activity*. (b) Heatmap of gene expression for *voltage-gated calcium channel activity* genes in iN control and iN deletion cells. (c) Bar plot of gene expression of *voltage-gated calcium channel activity* genes in fibroblast and iN cells showing the result would not have been found in a non-neuronal cell-type.

**Figure S7:**
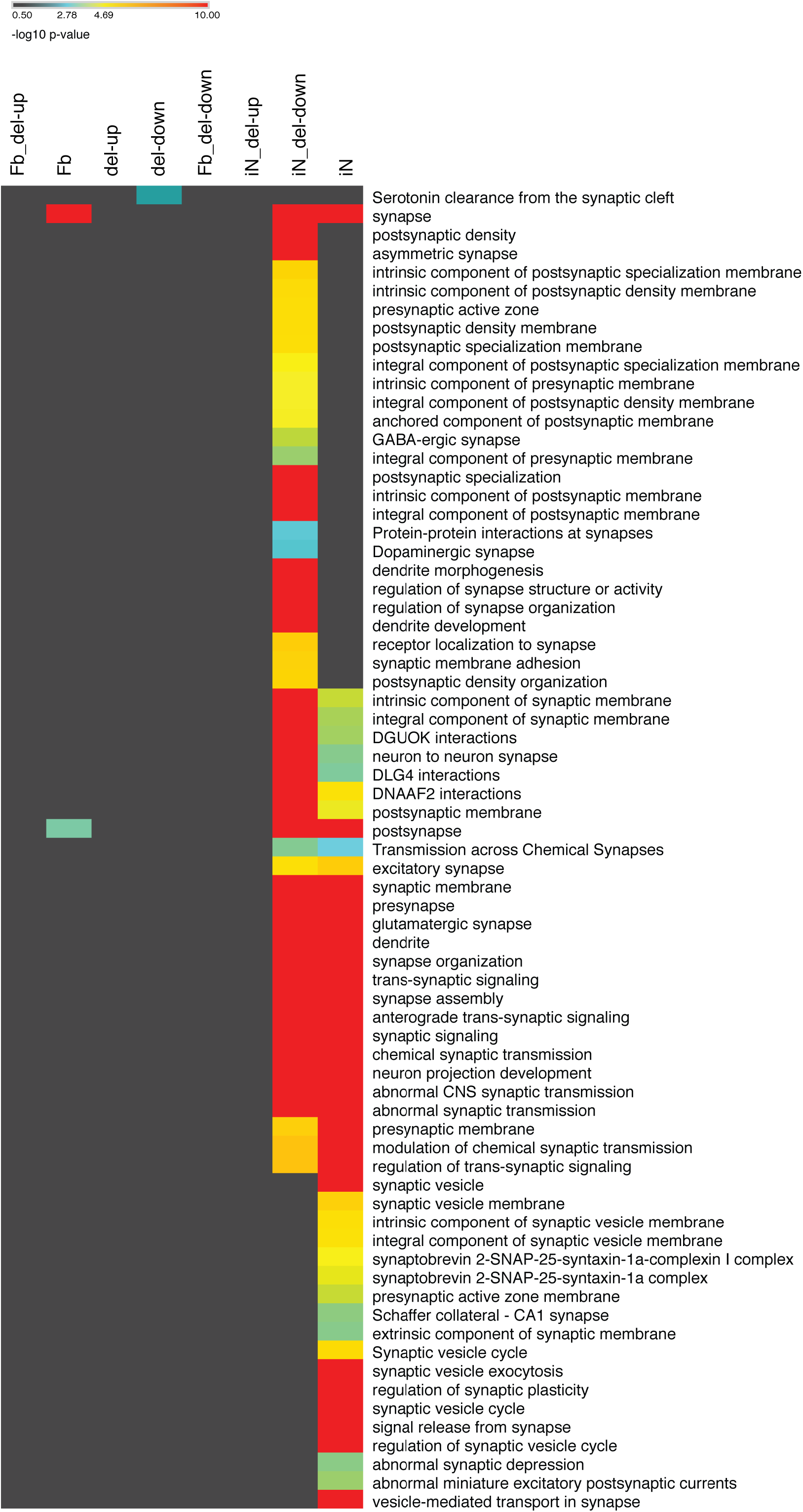
Comparison of synaptic pathways in iN and iN_del-down modules. Larger view of main Figure 6: p-values for synaptic pathways enriched in iN module or iN_del-down module showing pathways unique to downregulation in 22q11del iNs and not iN cells in general. Color code indicates −log10 pvalue.

**Figure S8:**
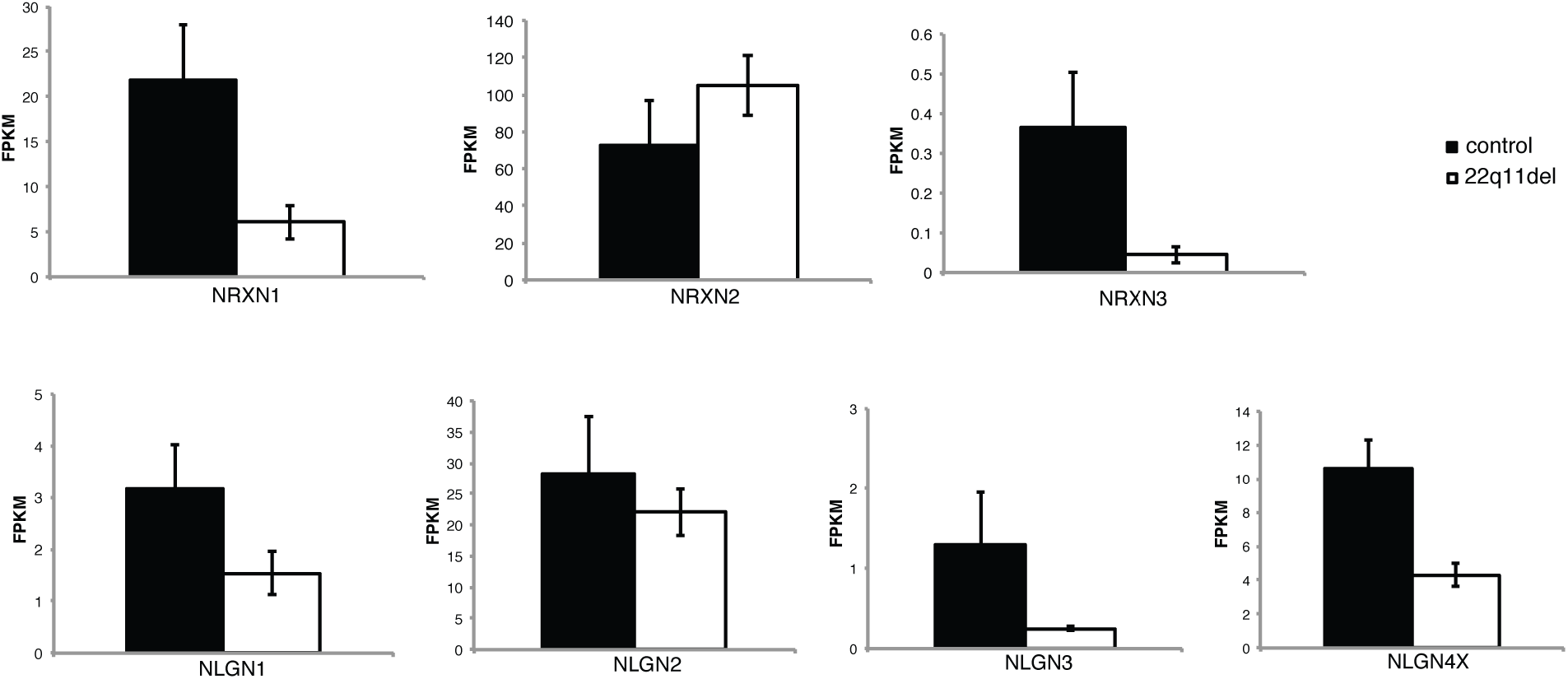
RNA expression of neurexin and neuroligin genes. Bar plots for expression of neurexins and neuroligins in iN control and deletion cells.

